# The complete, functional and dynamic cycle of the bacterial Initiation Factor 3

**DOI:** 10.1101/579326

**Authors:** Jose A. Nakamoto, Roberto Spurio, Andrey L. Konevega, Attilio Fabbretti, Pohl Milón

**Author notes:** Corresponding Author: Centre for Research and Innovation, UPC-Villa, Avenida Alameda San Marcos cuadra 2, Chorrillos, 15067 Lima, Peru.

## Abstract

Initiation factor 3 (IF3) is an essential protein that enhances the fidelity and speed of bacterial initiation of mRNA translation. The dynamic interplay between the two independent IF3 domains, their alternative binding sites, and the mechanism that ensures translation initiation fidelity remains elusive. Here, we show that the functional positioning of IF3 domains occurs at velocities ranging over two orders of magnitude, driven by each 30S initiation ligand. IF1 and IF2 rapidly promote the accommodation of IF3 on the 30S platform with the C-terminal domain moving towards the P site. Reversion of this movement is triggered by decoding the mRNA start codon and rate limits translation initiation. Binding of the tRNA results in the concomitant accommodation of the N-terminal domain of IF3, largely dependent on the mRNA and initiator tRNA. 70S initiation complex formation promotes the closing and dissociation of IF3, recycling the factor for a new round of translation initiation. Altogether our results unveil the kinetic spectrum of IF3 conformations and highlight fundamental movements of the factor that ensure accurate translation initiation.

## Introduction

IF3 is an essential bacterial protein, consisting of two domains (IF3C and IF3N) separated by a hydrophilic, lysine-rich, linker [1–3]. IF3 is involved in all steps of the bacterial translation initiation driving the 30S ribosomal subunit trough the transition from the 30S initiation complex (30S IC) to a productive 70S initiation complex (70S IC) [4–6]. When bound to the 30S subunit, IF3 prevents the premature 50S subunit association and increases the rate of the P site codon-anticodon interaction between fMet-tRNA^fMet^ and the initiation triplet of the mRNA [5–8]. As a direct consequence, IF3 acts as an initiation fidelity factor by increasing the dissociation rate of non-canonical and pseudo-30S initiation complexes [8–10].

Although IF3 functions during the initiation phase of protein synthesis are well established, dynamic aspects of IF3 binding on ribosomal subunits and the release of the factor after the formation of productive 30S IC are still debated [11,12]. A model where IF3C and IF3N domains independently bind the 30S ribosomal subunit was initially proposed in the early ‘80s [13] and then confirmed by NMR analysis and time-resolved chemical probing experiments [14,15]. Several studies have dealt with the assignment of a topographical localization of IF3 on the 30S ribosomal subunit, producing conflicting conclusions [16–19]. Recently, CryoEM experiments demonstrated that IF3 undergoes large conformational changes to facilitate the accommodation of the initiator tRNA (fMet-tRNA^fMet^) into the P site for start codon recognition [20]. These experiments, together with other recent structural reconstructions, contributed to the identification of a minimum of four different IF3 arrangements on the 30S ribosomal subunit and two alternative binding sites for each domain of the factor [20–22]. IF3C can interact with two distinct binding sites on the 30S subunit, either at the P site and in contact with IF1 (C2 position in Fig 1), or at helix 45 and helix 24, away from both IF1 and the P site (C1 position in Fig 1). Similarly, IF3N can occupy two binding sites, either on the 30S platform near S11, or on the elbow of tRNA^fMet^, herein called NR and NT, respectively. Despite these detailed structural data, a dynamic correlation between IF3 interaction sites, the ligands present on the 30S subunit and the functions of the factor remained elusive.

**Figure 1.**
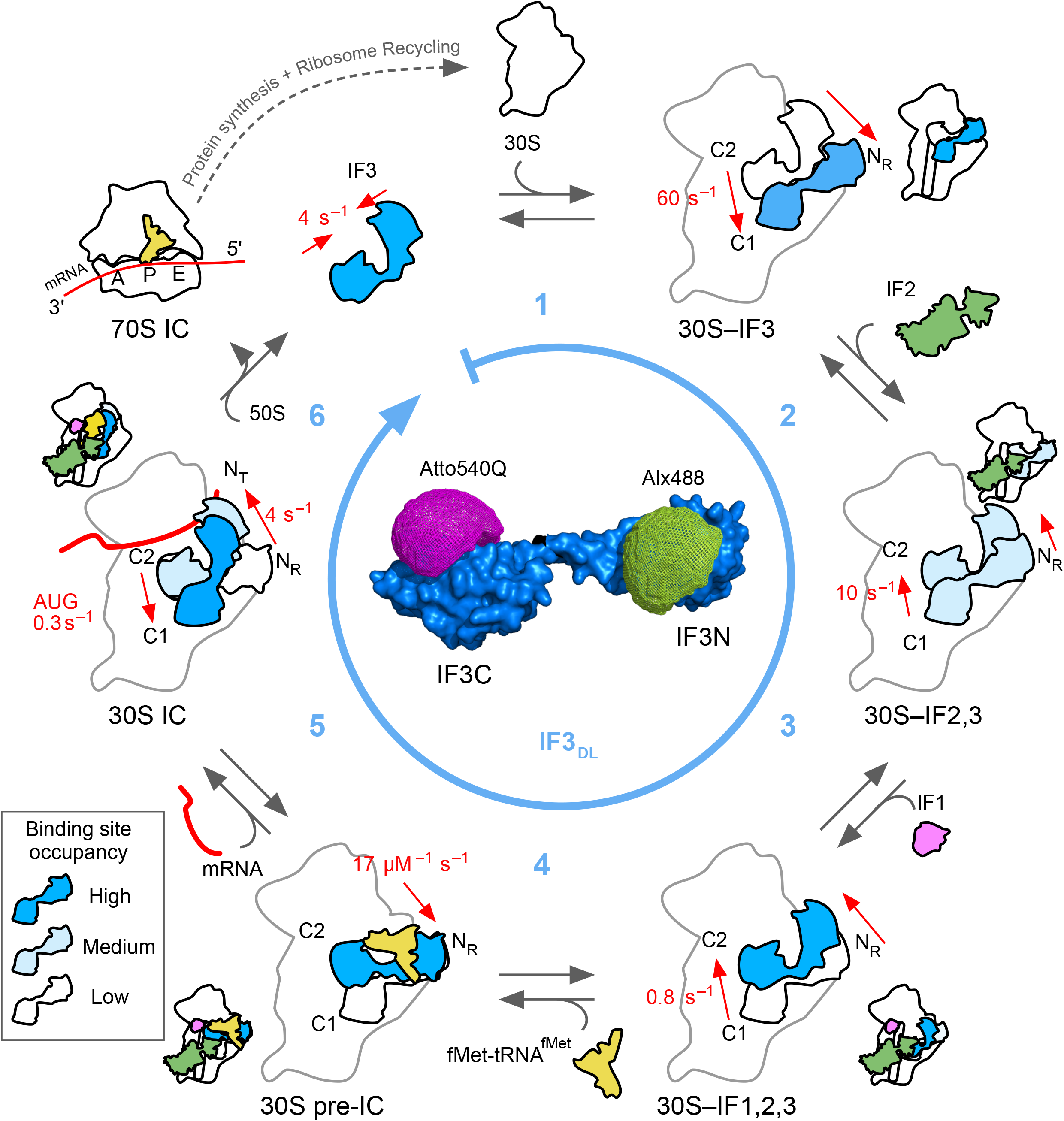
Scheme of IF3 motions during bacterial translation initiation. Conformational accommodations of IF3 along the full cycle of translation initiation was monitored by intramolecular FRET between dyes specifically linked to each domain of the factor (IF3_DL_, inset) and structural modeling of IF3_DL_ on available 30S IC structures [20]. Round arrow (sky blue) indicates the direction of complex progression by measuring IF3_DL_ accommodation as a function of the 30S subunit (white), IF2 (green), IF1 (pink), fMet-tRNA^fMet^ (yellow), the mRNA (red), and the 50S subunit. Sky blue numbers indicate the main reactions studied here. Red arrows and values indicate the directionality and rates of IF3 movements measured here. Alternative binding sites for IF3N are: NR (30S-bound) and NT (fMet-tRNA^fMet^-bound). Alternative binding sites for IF3C are indicated as C1 or C2. 30S pre-IC indicates the pre-initiation complex while mRNA is indicated by a red line in the 30S and 70S ICs.

In this study, we use a combination of Föster Resonance Energy Transfer (FRET), presteady state kinetics techniques and molecular modeling to monitor IF3 movements on the ribosome and to construct a comprehensive model depicting the dynamics of IF3 domains along the full cycle of translation initiation, from the initial binding until the recycling of the factor (Fig 1).

## Results and Discussion

To monitor the conformational rearrangements of IF3, we used a fluorescent derivative of the factor harboring donor and silent acceptor dyes specifically linked to the N and C terminal domains, respectively (Fig1, inset) [23]. Double-labeled IF3 (IF3_DL_) allows to monitor intramolecular FRET changes resulting from the movement of IF3 domains with respect to each other. Donor fluorescence values were used to measure FRET changes of IF3_DL_ in a stopped-flow apparatus, allowing to monitor IF3 movements in real-time. An increase in time of donor fluorescence indicates that IF3 domains are moving apart and, *vice versa*, a decrease of the observed fluorescence indicates that the domains are getting close to each other (Supp. Fig 1) [23]. Binding of IF3_DL_ to the 30S ribosomal subunit resulted in an increase of fluorescence, consistent with the factor transiting from a closed conformation in solution to a more opened one while bound to the 30S subunit (Fig 2A). A biphasic dependence over time was observed with both apparent rates saturating with 30S concentration (Fig 2B, Supp. Fig 2A). Kinetic analysis of the 30S concentration dependence indicated that after the initial binding (10^3^ μM^−1^ s^−1^, [4]), IF3 stretches at an average velocity of 60 s^−1^. Our data is in agreement with time resolved chemical probing and NMR spectroscopy analysis showing that IF3C and IF3N sequentially interact with the 30S ribosome [14,15].

**Figure 2.**
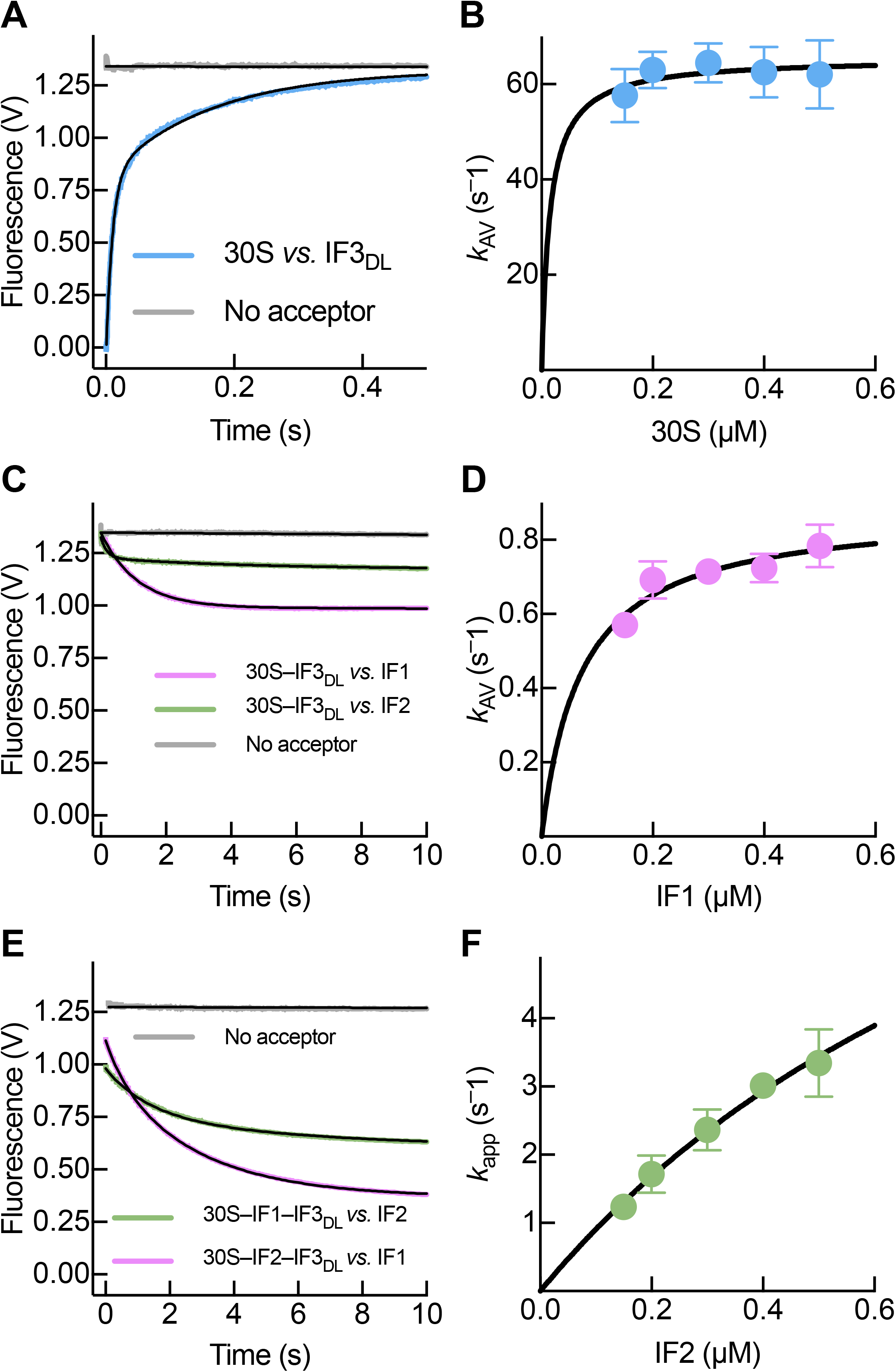
IF1 and IF2 rapidly promote a compact layout of IF3 on the 30S platform. Stopped-flow analysis of IF3_DL_ conformational transitions during binding to the 30S subunit and binding of IF1 and IF2 to the 30S–IF3_DL_ complex. A Time course of donor fluorescence change of 0.04 μM IF3_DL_ during binding to 0.12 μM 30S (sky blue). B IF3_DL_ binding velocities as a function of 30S concentration. Averaged rates were used to estimate the overall velocity of IF3 opening (Equation 4). C Time courses of 0.5 μM IF1 (pink) or IF2 (green) binding to 0.05 μM 30S–IF3_DL_ complexes. D IF1 concentration dependence of the averaged apparent rates of IF3_DL_ closure upon binding of IF1 to the 30S–IF3_DL_–IF2 complexes. E Time traces of IF3_DL_ donor fluorescence change as function of 0.5 μM IF1 (pink) or IF2 (green) binding to 0.05 μM 30S–IF3_DL_–IF2 or 30S–IF3_DL_–IF1 complexes, respectively. F IF2 concentration dependence of the apparent rates of IF3_DL_ closure upon binding of IF2 to the 30S subunit. Grey lines in (A), (C), and (E) represent reactions with IF3 labelled only with the donor fluorophore at IF3N, without the quencher molecule at IF3C (no acceptor) to verify that the fluorescence signal of IF3_DL_ depended on distance changes between fluorophores. Continuous lines represent non-linear fitting with exponential (A, C, E, Equations 1-2) or hyperbolic (B, D, F, Equation 3) functions. Each time trace results from the average of 7-10 independent measurements. Error bars in (B), (D), and (F) represent standard deviations.

To position each domain of IF3_DL_ on the 30S subunit, we modeled the dyes on each IF3 domain at all possible binding combinations from recent 3D reconstructions [20] (Supp. Fig 3 and 4). The resulting accessible volumes for the dyes allowed to calculate the theoretical distances between fluorophores and domains (Supp. Fig 3). The structural modeling of IF3_DL_ combined with FRET changes as a function of 30S concentration suggests that in the absence of any other ligand, IF3 acquires the most extended layout with IF3C occupying the C1 site and IF3N moving into the N_R_ site with an interdomain distance of about 60 Å (Fig 1, step 1, Supp. Table 1). This IF3 layout is affected by all ligands of the 30S subunit; however, none promotes an IF3 opening as extended as that in the 30S–IF3_DL_ complex (see below).

Binding of IF1 to the 30S–IF3_DL_ complex resulted in a decrease of fluorescence over time, indicating that IF3 domains transit from an extended to a more compact layout on the 30S subunit (Fig 2C) in agreement with single molecule analysis [24]. Analysis of FRET time traces obtained at increasing concentrations of IF1 indicated that IF3_DL_ closes at an average speed of 0.8 s^−1^ (Fig 2D, Supp. Fig 2C). Structural modeling of IF3_DL_ on the 30S subunit suggests that IF1 enhances the movement of IF3C from the C1 to the C2 binding site (Fig 1, step 3), partially accounting for the observed reduction of interdomain distances. The extent of fluorescence reduction suggests that also IF3N is displaced towards IF3C. Binding of IF2 also promoted a closure of IF3 at over 10-fold faster velocity (10 s^−1^) than those observed for IF1 (Fig 1, step 2, Fig 2F); however, with a different amplitude dependencies on the concentration of the factor (Fig 2C, Supp. Fig 2B). IF1 and IF2 cooperatively maximized the extent of IF3 closing (Fig 2E). Altogether, our data indicates that the intermediate 30S–IFs complex promotes an IF3 layout where IF3N dissociates from the NR binding site and IF3C moves towards the C2, near the P site. While occupation of the C2 binding site would interfere with the formation of inter-subunit bridges (reviewed in [25]), positioning of IF3N would contribute to shape the tRNA^fMet^ binding site.

Thus, the 30S–IFs complex assembles in ≈ 30 ms [4] and rearranges in ≈ 1 s (this work). This intermediate of translation initiation is characterized by a high affinity of IF3, and consequently, it prevents the binding of the 50S subunit [4] [26]. On the other hand, when a canonical start codon of the mRNA is present in the P site, the initial interaction of fMet-tRNA^fMet^ with the 30S complex is followed by codon-anticodon formation. The resulting 30S IC, in comparison to the pre-IC, displays an increased dissociation rate constant (*k*_off_) of IF3 from the 30S subunit [4,5]. To probe how IF3 interdomain dynamics relate to the differential off rates of the factor, we first measured fMet-tRNA^fMet^ binding to 30S complexes containing IFs but lacking the mRNA. The resulting fluorescence traces increased with time, indicating a re-accommodation of either (or both) IF3 domains, moving away from each other (Fig 3A). Kinetic analysis obtained at increasing concentrations of fMet-tRNA^fMet^ showed a linear dependence of velocities with tRNA concentrations, consistent with IF3_DL_ monitoring the initial interaction of the fMet-tRNA^fMet^ with 30S pre-IC lacking the mRNA (Fig 3B). Our results indicate that the fMet-tRNA^fMet^ rapidly binds to the 30S complex, *k_1_* = 17 μM^−1^ s^−1^, and can readily dissociate, *k_−1_* = 1 s^−1^. Both kinetic parameters are in agreement with previous studies where different FRET pairs were used to monitor fMet-tRNA^fMet^ binding to 30S pre-ICs [27]. Thus, the initial interaction of fMet-tRNA^fMet^ entails the partial opening of IF3, probably as a consequence of displacing IF3N rather than IF3C. Altogether, the formation of the 30S pre-IC lacking the mRNA entails an IF3 layout where IF3C is positioned at the C2 site and IF3N is pushed by the initiator tRNA towards the N_R_ site. The reaction is rapid and under our experimental conditions (1 μM of fMet-tRNA^fMet^) IF3 accommodation during 30S pre-IC formation shall take less than 50 ms.

**Figure 3.**
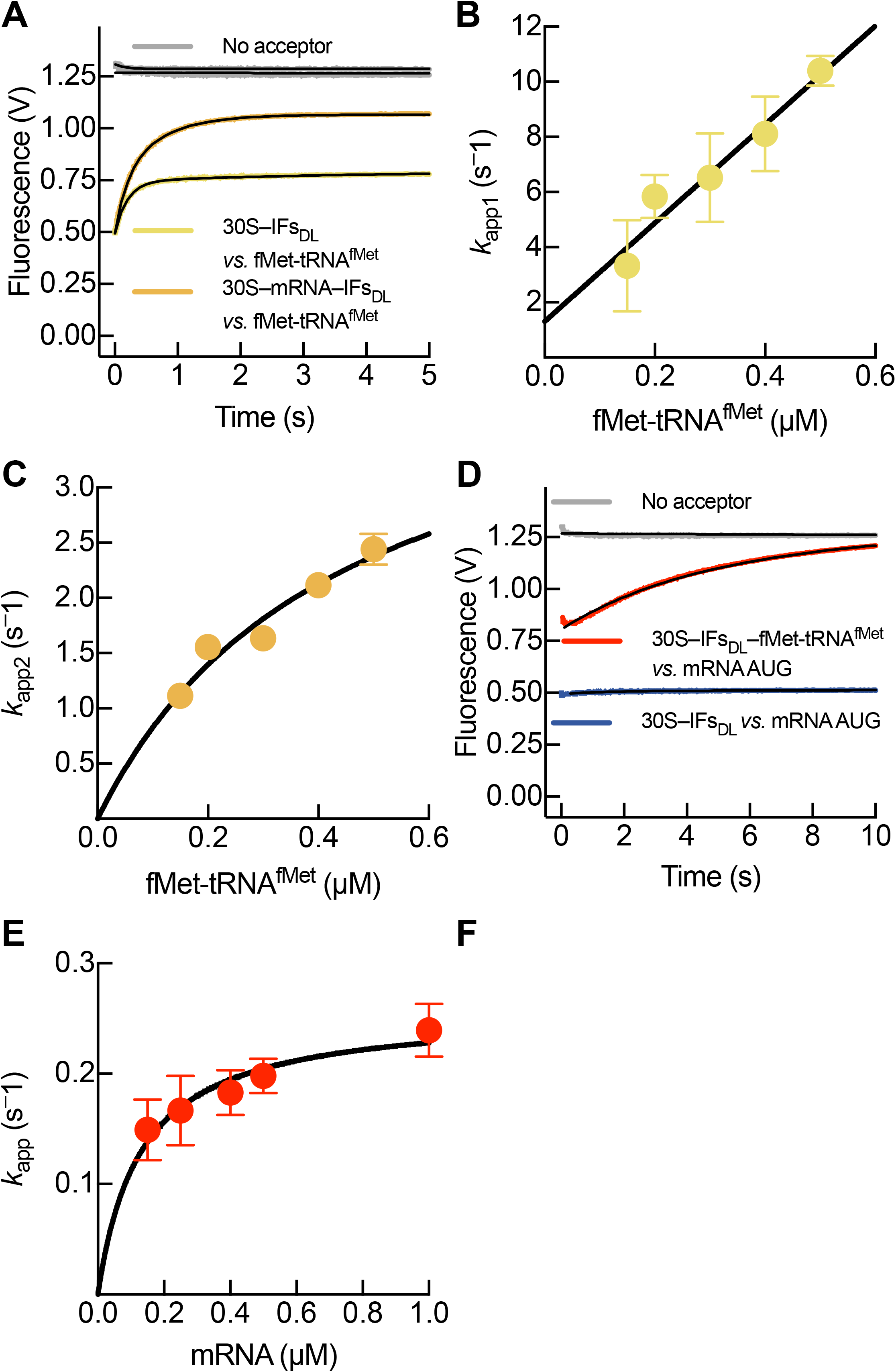
fMet-tRNA^fMet^ and decoding of the mRNA start site promote an open layout of IF3 on the 30S platform. A Time course of donor fluorescence change of IF3_DL_ during binding of 0.5 μM fMet-tRNA^fMet^ to 0.05 μM 30S pre-IC in the presence (orange) or absence (yellow) of the mRNA. B, C fMet-tRNA^fMet^ concentration dependence of the first (B) and second (C) apparent rates of IF3_DL_ opening upon binding of the initiator tRNA to 30S pre-ICs. D Time courses of donor fluorescence change of IF3_DL_ during binding of 0.5 μM mRNA to 0.05 μM 30S pre-IC in the presence (red) or absence (blue) of fMet-tRNA^fMet^. E mRNA concentration dependence of the apparent rates of IF3_DL_ closure upon binding to the 30S pre-ICs containing fMet-tRNA^fMet^. The control curves, indicated by grey lines in (A) and (D), represent reactions carried out in the presence of IF3 lacking the acceptor at the C-terminus subunit (no acceptor) to assess whether the fluorescence signal was dependent on the distance change between fluorophores. Continuous lines represent the fitting with exponential (A) and (D) (Equations 1-2), linear (B) or hyperbolic (C) (E) (Equation 3) functions. Each time trace results from the average of 7-10 independent measurement. Error bars in (B), (C), and (E) represent standard deviations.

The extent of IF3_DL_ signal as a function of fMet-tRNA^fMet^ recruitment depended on the presence of the mRNA. As indicated in Fig 3A, mRNA-programmed 30S complexes produced larger amplitudes of IF3_DL_ opening if compared with complexes without mRNA. In addition, the fluorescence trace appeared biphasic in time, indicating a further accommodation of IF3_DL_ following the initial binding of fMet-tRNA^fMet^ (Fig 1, step 4, Fig 3A). Kinetic analysis of time courses obtained at increasing concentrations of fMet-tRNA^fMet^ showed that the velocities of the fMet-tRNA^fMet^ accommodation step in 30S preICs containing the mRNA tended to saturate at 4 s^−1^ (Fig 3C, Supp. Fig 2D). These experimental data, coupled with the structural modeling, support a model where IF3 accommodates the fMet-tRNA^fMet^ on the P site, most likely through IF3N evaluating intermediates of initiator tRNA and IF3C moving back towards the C1 position (Fig 1, steps 4,5). Both reactions push IF3 domains away from each other.

However, the complexity of the above reactions precluded us to obtain accurate information on the kinetics of mRNA start site decoding. To overcome this limitation, we set up an experiment where 30S pre-ICs were pre-bound to fMet-tRNA^fMet^ and mixed with increasing concentrations of mRNA. Mixing of the mRNA with 30S pre-ICs resulted in an increase of IF3_DL_ fluorescence, indicating an increase of distance between IF3 domains with velocities saturating at 0.3 s^−1^ (Fig 3D,E, Supp. Fig 2E). The increase of fluorescence observed in this experimental setup is likely the result of the displacement of IF3C from C2 towards the C1 binding site (Fig 1, step 5) rather than repositioning IF3N, which is affected by the presence of the pre-bound initiator tRNA (see above). Accordingly, structural modeling of IF3_DL_ on 30S IC structures indicate that the IF3C displacement accounts for 10 Å change of distance between the fluorophores on IF3 domains (Supp. Fig 3 and 4). A comparison of the velocities for all IF3 movements suggests that the displacement of IF3C towards the C1 binding site is the slowest, likely occurring during decoding of the mRNA start site. Thus, our results show that the IF3C displacement is the slowest step, rate-limiting the progression of 30S IC formation and taking about 3 s.

The progression towards 30S IC formation entails an isomerization or locking step between the 30S pre–IC and 30S IC (reviewed in [28]). Both complexes are identical in terms of bound ligands; yet, they differ in their biochemical properties. While the off rates of IF1, IF2, mRNA, and initiator tRNA are greatly reduced in the locked 30S IC, that of IF3 is increased [4]. Thus, the alternative binding sites of IF3C can be associated to the differential IF3 off rates observed in the unlocked and locked 30S complexes. While the C1 binding site would represent fast off rates, the C2 can be responsible of the slow dissociation rates for IF3. The C1 binding site can be proposed as both, the entry and exit position for IF3C, while the C2 site can be proposed as the active site for IF3C.

Joining of the 50S to 30S ICs results in an unstable 70S pre-IC which upon release of IFs allows the formation of a 70S IC capable of translating the in-frame mRNA [5,11]. The reaction appears to be sequential [11] and to be tightly regulated by IF3 [5,26,29]. 50S joining to 30S ICs with IF3_DL_ resulted in a fluorescence decrease over time, indicating a closure of the factor upon dissociation (Fig 1, step 6, Fig 4A). Kinetic analysis of time courses at increasing concentrations of 50S suggests that the overall rate of dissociation and subsequent closure of IF3_DL_ occurs at 4 s^−1^, a rate very similar to previous reports where IF3 dissociation was measured with FRET between the factor and fMet-tRNA^fMet^ (Fig 4B, Supp. Fig 2F) [5,11].

**Figure 4.**
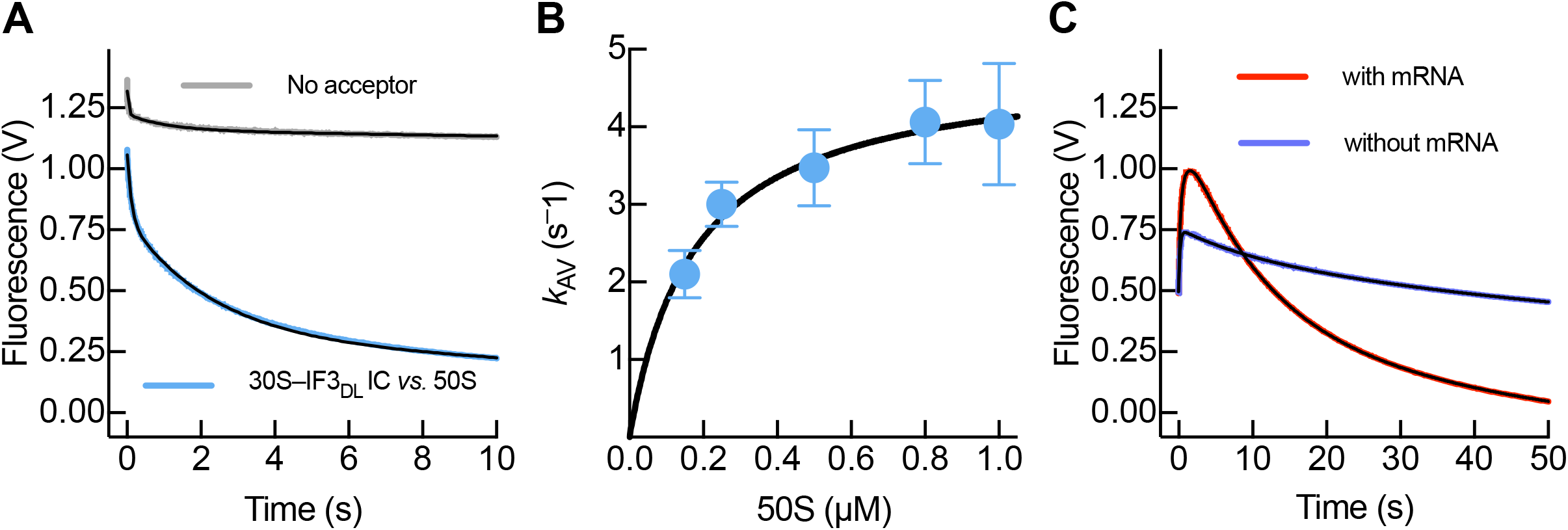
70S IC formation is accompanied by IF3 closing during dissociation. A Time course of donor fluorescence change of IF3_DL_ during binding of 0.5 μM 50S to 0.05 μM 30S IC (sky blue). Grey line indicates a control using an IF3 without the acceptor dye (as in Fig 2 and 3). B 50S concentration dependence of the average rates of IF3_DL_ closure upon binding of the large subunit. Error bars represent standard deviations. C Time courses of donor fluorescence change of IF3_DL_ during the concomitant binding of 0.5 μM 50S and fMet-tRNA^fMet^ to 0.05 μM 30S pre-IC in the presence (red) or absence (sky blue) of the mRNA. Continuous lines represent the fitting with exponential (A) and (C), or hyperbolic (B) functions. Each time trace results from the average of 7-10 independent measurements. Error bars in (B) represent standard deviations.

Previous reports showed that the efficiency and velocity of 70S IC formation (i.e. subunit joining, GTP hydrolysis, Pi release, and IFs dissociation) strictly depended on the tRNA and programmed mRNA [5,6,11,26,30]. When we measured the concomitant arrival of fMet-tRNA^fMet^ and 50S to 30S pre-ICs with and without mRNA, we could observe a time dependence of IF3_DL_ fluorescence change indicating the opening of the factor due to initiator tRNA binding (increase from 0.5 to 1.0) and the following closing due to IF3 ejection from the resulting 70S IC (decrease from 1.0 to 0.0) (Fig 4C). The rate of the latter was very similar to that monitored upon mRNA binding to 30S pre-ICs, indicating that decoding of the start codon and the subsequent movement of IF3C to the C1 site, rate limits 70S IC formation. In the absence of the mRNA, IF3_DL_ in the 30S pre-IC still sensed the arrival of fMet-tRNA^fMet^; however, IF3 dissociation was missing and the factor appeared to adopt its initial layout (Fig 4C). Whether initiator tRNA stays bound or dissociates from these complexes, remains an open question.

Our intramolecular FRET approach to measure IF3 interdomain layouts during 30S IC formation and comparison with theoretical distance changes between fluorophores revealed a timely model for the functional IF3 cycle: from binding to dissociation (Fig 5). The combination of both approaches indicates subsequent movements of IF3 domains, in response to the recruitment of each 30S ligand and within a detailed temporal framework. While IF1 and IF2 cooperatively allow IF3C sensing start codon decoding, movements of IF3N appear to be coupled to fMet-RNA^fMet^ binding and subsequent accommodation. Our study reveals the kinetics associated to each movement of the factor with remarkable insights. First, IF3C moves to the C2 site in 0.1 - 1 s, induced by IF1 and IF2 recruitment, (Fig 1, Steps 2,3) likely responsible for preventing the premature joining of the 50S subunit. Second, IF3N dissociates from the 30S platform to interact with fMet-RNA^fMet^, with a subsequent tRNA accommodation towards the P site (Fig 1, Step 4,5). Third, decoding of the mRNA start site by the P-site initiator tRNA displaces IF3C towards the C1 site. This event, which is a rate-limiting step for the formation of the 30S IC (≈ 3 s) allows joining of the 50S subunit (Fig 1, Step 5). Finally, IF3 dissociates during 70S IC formation in less than 250 ms and the factor is now ready to bind a newly available 30S subunit (Fig 1, Step 6, Fig 5).

**Figure 5:**
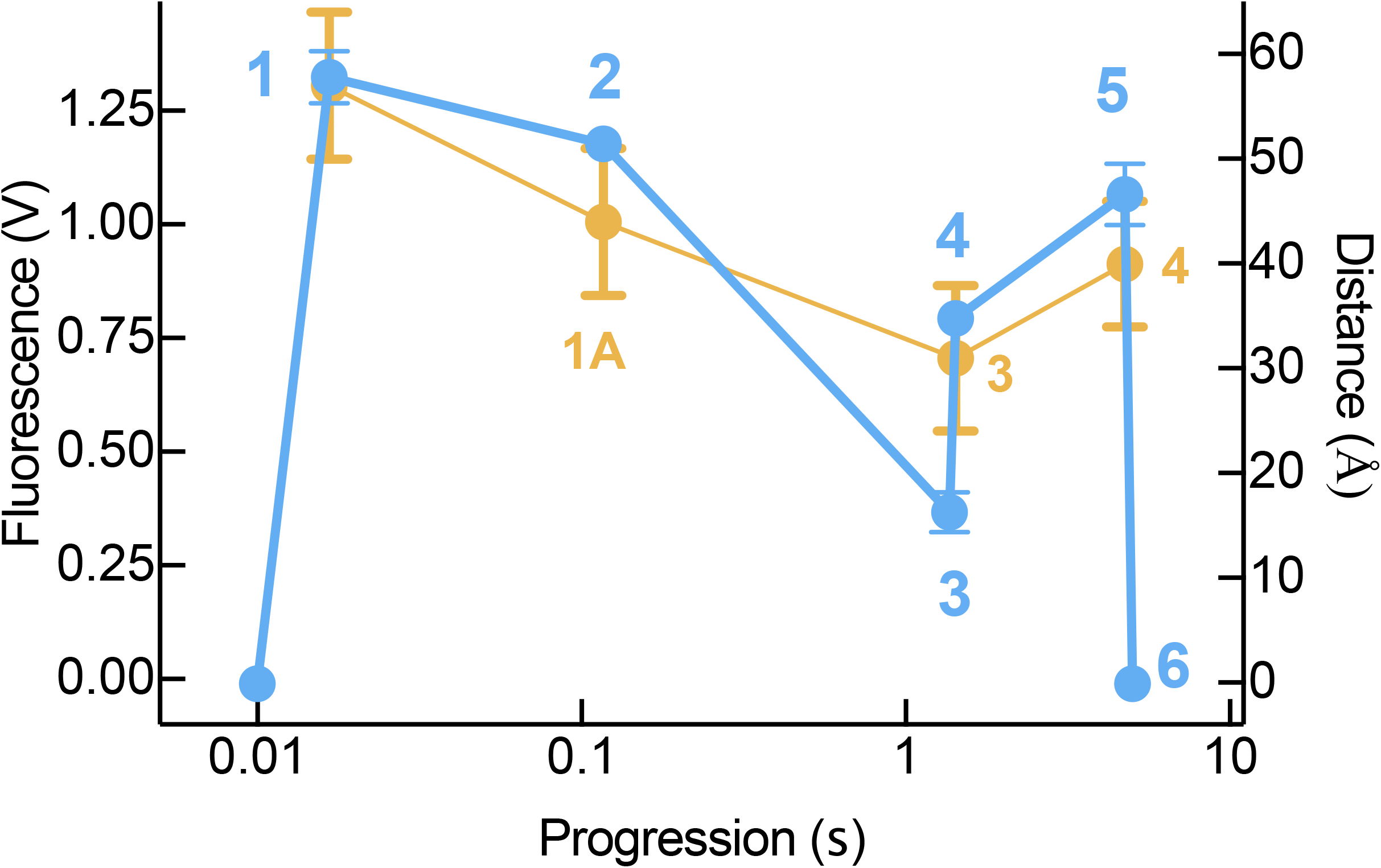
Intramolecular distances between IF3 domains during progression of translation initiation. Comparison between the amplitude changes of IF3_DL_ for all the reactions measured here (sky blue, left axis) and distances between the fluorophores on the corresponding complexes as analyzed by molecular modeling (orange, right axis). Sky blue numbers indicate the reactions measured here (refer to Fig 1). Orange numbering refer to the 30S PICs 1A, 3, and 4 complexes from [20]. *x*-axis represent the progression time of each reaction as calculated from the saturating rates measured here. Error bars represent standard deviations from the total amplitudes for each reaction (sky blue) or from the distance distribution of IF3_DL_ fluorophores calculated by structural modelling of the complexes mentioned above (orange, Supp. Figure 3 and 4, Supp. Table 1).

A model where each IF3 domain moves independently can fully accommodate our current understanding of the dynamics of the factor, on and off the ribosome (movie in supplementary materials). In this perspective we report the dynamics and directionality of IF3 movements resulting from the interaction of each translation initiation ligand with the 30S subunit. We observe the sequential reduction of inter-domain distance towards the assembling of the 30S pre-IC lacking the initiator tRNA. Although, from our FRET studies we cannot derive the position of IF3 domains on the ribosome, we observe an interdomain distance trend similar to the hypothetic progression of available 30S IC structures and modeling of IF3_DL_ (Fig 5, steps 1 to 3) [20]. The reduction of IF3 interdomain distances appears more evident in the FRET data (step 3, sky blue) if compared to that derived from modelling of the corresponding structures (Fig 5, complex 3 (orange)). This difference is likely to arise from the C2 to C1 movement of IF3C, or from the concomitant movement of IF3N. The latter is an appealing model as it would entail a progressive induced fit of the fMet-tRNA^fMet^ binding architecture. In this context, IF3C moving to the C2 site would be coupled to the dissociation of IF3N from the N_R_ site to adopt a layout that is ready to interact with the initiator tRNA. This alternative model is supported by single molecule studies where interdomain distances were shown to be influenced by IF1 and IF2 [24]. In this context, binding of fMet-tRNA^fMet^ and decoding of the mRNA start site would uncouple IF3 movements, promoting IF3C to move towards the C1 position while the IF3N remains tRNA-bound.

In a cellular context, the progression of translation initiation would result in at least two kinetically stable intermediate complexes, 30S–IFs and the 30S pre-IC. The former would ensemble in around 1 s, being able to reject the 50S by maximal occupancy of the C2 site by IF3C. On the other hand, the 30S–IFs complex can rapidly recruit the initiator tRNA at the P site by providing a functional binding pocket with IF3C at the C2 site and IF3N ready to interact with the tRNA elbow. Coordinately, IF2 through its Carboxyl-terminal domain interacts with the formyl group of the initiator tRNA. The transition towards a 30S pre-IC, harboring all ligands, would be limited by the availability of mRNA translation initiation regions (TIRs). Finally, upon decoding of the mRNA start site, the displacement of IF3C from the C2 site induces 30S IC locking, limiting the overall progression rate of translation initiation. This step would allow to build 70S pre-ICs in less than 250 ms followed by a rapid and directional cascade of reactions that ultimately leads to a productive and ready-to-elongate 70S IC [11]

In summary, we describe the functional conformations of IF3 defining with high accuracy the distances and temporal framework (movie in supplementary materials) covered by both domains of the factor in correlation with all events taking place during the translation initiation process. Altogether, this work allows to accurately assign the speed and nature of elemental movements of IF3 behind its essential fidelity function.

## Materials and Methods

*Escherichia coli strains, expression vectors, cell growth and protein expression* Expression vectors pET24c containing *InfA*, *InfB*, *InfC*, and *InfC-E166C* were acquired commercially (GenScript, USA). Competent *E. coli* BL21 (DE3) cells were CaCl_2_ transformed (Mix & Go, ZymoResearch, USA) with expression vectors coding for IF1 *wt*, IF2 with an NTD His-tag, IF3 *wt* or IF3_E166C_. Typically, 2 L of Luria Bertani (LB) medium were used to grow the BL21 (DE3) strains to an OD_600 nm_ of 0.5. Protein expression was induced by adding 1 mM Isopropyl β-D-1-thiogalactopyranoside (IPTG, ThermoScientific, USA). Cells were allowed to express the factors for 3 hours prior to harvesting by centrifugation at 5000 x g at 4°C. Cells were lysed by sonication (20 cycles of 10 s followed by 30 s without sonication) in Buffer A (50 mM Hepes (pH 7.1), 100 mM NH_4_Cl, 10 mM MgCl_2_, 10 % Glycerol, 6 mM 2-mercaptoetanol) for IF1 and IF3; or Buffer B (50 mM Sodium Phosphate buffer (pH 7.7), 300 mM NaCl, 30 mM Imidazole) for IF2. Debris and supernatant were separated by centrifugation at 15,000 x g for 30 min a 4 °C.

### Purification of Initiation Factors

Initiation factors IF1 and IF3 were purified by Cation exchange chromatography on a HiTrap™ SP HP column (GE Healthcare Life Sciences, Uppsala, Sweden). Cleared lysates were loaded onto the column (1 mL column volume, CV) and subsequently subjected to a linear NH_4_Cl gradient (0.05 – 1 M) in Buffer A using a HPLC system (Jasco, Japan). IF3 and IF1 eluted at 700 mM and 400 mM NH_4_Cl, respectively. Best separation conditions were found at 1 mL/min flow rate and 20 CVs long gradient, collecting fractions of 1 mL each. Protein elution was followed by absorbance at 290 nm and SDS-Polyacrylamide Gel Electrophoresis (SDS-PAGE, 15 %). While IF3 eluted with an elevated degree of purity, IF1 fractions contained high molecular weight contaminants. Protein contaminants were eliminated by subjecting the combined IF1 fractions to Amicon^®^ Ultra 30K Da centrifugal filters (Merck, Germany) followed by concentration on a HiTrap™ SP HP (GE Healthcare Life Sciences, Uppsala, Sweden), single step eluted with 1 M NH_4_Cl in Buffer A. Eluted proteins were dialyzed against Storage buffer (50 mM Tris-HCl (pH 7.1), 200 mM NH_4_Cl, 10 % Glycerol, 6 mM 2-mercaptoethanol) and small aliquots were stored at – 80 °C. Purity was assayed by 15% SDS-PAGE and total protein staining using blue coomassie.

Initiation factor IF2 was purified by affinity chromatography on a Histrap FF crude column (1 mL). Supernatants were manually loaded to the column, followed by 5 mL of wash buffer (50 mM Imidazole in Buffer B) and to 2.5 mL of elution buffer (250 mM Imidazole in Buffer B). The first 1.5 mL were collected and subjected to dialysis in Buffer A. Finally, IF2 was further purified by Cation exchange chromatography as explained above for IF1 and IF3.

### Double labeling of IF3

IF3_E166C_ was subjected to extensive dialysis in Labeling buffer (50 mM Hepes (pH 7.1), 100 mM NH_4_Cl, 10 % Glycerol and 0.5 mM TCEP) in a D-Tube™ Dialyzer Maxi (EMD Millipore Corp, Billerica, USA) to remove traces of 2-mercaptoethanol as the reducing agent strongly inhibits the coupling to maleimide linked dyes to cysteines. First, the C-terminal was labeled at the recombinant cysteine (C166) as it is exposed and efficiently reacts with maleimide derivatives. A 10-fold excess of Atto-540Q maleimide (Atto-Tec GmbH, Siegen, Germay) over IF3_E166C_ was incubated in Labelling buffer for 20 minutes. The reaction was stopped by the addition of 6 mM 2-mercaptoethanol. The modified IF3_C540Q_ was purified from unreacted dyes on a HiTrap SP HP column as described above. After 10 CV washes with Buffer A containing 100 mM NH_4_Cl, a single step elution was applied using 3 mL of 1 M NH_4_Cl in Buffer A. Typically, full protein recovery is achieved in 0.5 mL and elution of the labeled protein is readily visible. IF3_C540Q_ was subsequently dialyzed as mentioned above in labeling buffer containing 2 M UREA. The partial denaturation of IF3 results in the exposure of the otherwise buried cysteine at position 65 of the NTD. The unfolded protein was incubated with a 10-fold molar excess of Alexa488 maleimide (Invitrogen, USA) for 1h at RT, mild shacking was applied. IF3_C540Q–N488_ (IF3_DL_) was purified from the unreacted dye as described above using HiTrap SP HP column. Eluted proteins were dialyzed against Storage buffer and small aliquots were stored at – 80°C. Purity and efficiency of labeling was assayed by 15% SDS-PAGE where fluorescence was observed under a UV trans-illuminator and total protein was determined by Blue Coomassie staining.

### Ribosomal subunits, fMet-tRNA^fMet^ and mRNAs

30S and 50S ribosomal subunits were purified from tight coupled 70S ribosomes by sucrose gradient centrifugation under dissociative conditions (3.5 mM MgCl_2_), essentially as described [31]. 30S subunits were reactivated by 20 mM MgCl_2_ treatment for 30 min at 37 °C prior to use. fMet-tRNA^fMet^ was *in vitro* aminoacylated, formylated, and purified by reverse phase HPLC as described [31]. The model mRNA was chemically synthetized and commercially acquired from Trilink (USA) or Microsynth (Switzerland) with the following sequence: AAA CAA UUG GAG GAA UAA GGU AUG UUU GGC GGA AAA CGA.

### Stopped–Flow measurements and analysis

Fluorescence stopped-flow measurements were performed using a SF-300X stopped-flow apparatus (KintekCorp, USA) or a SX20 (Applied Photophysics, UK) by rapidly mixing equal volumes of each reacting solution. The excitation wavelength for Alexa 488 was 470 nm. The emission signal was measured after a long-pass optical filter with a 515 nm cut-off. 1000 points were acquired in each measurement. 7 to 10 replicates were recorded for each reaction and subsequently averaged. All stopped flow reactions were performed in TAKM10 buffer (50 mM Tris (pH 7.5), 70mM NH_4_Cl, 30 mM KCl, 10 mM MgCl_2_, 6 mM 2-Mercaptoethanol) at 25°C. Unless otherwise stated, 30S complexes were prepared by mixing 0.1 μM 30S, 0.08 μM IF3_DL_, 0.3 μM IF1,2, 100 μM GTP, 0.5 μM mRNA or fMet-tRNA^fMet^. Mixtures were incubated at 37 °C for 30 min and centrifugated at 14000 x g for 5 min prior to the measurement. Fluorescence time courses were baseline removed, and to highlight the overall conservation of the signal along the full cycle of IF3, an amplitude factor corresponding to the previous reaction was added. All graphical representations have the same y axis range and represent change of Volts as measured by the Instrument Photomultiplier. Time courses were analyzed by non-linear regression with one or two exponential terms (equations 1 and 2) using the Prism Software v. 7.0 (Graphpad, USA).

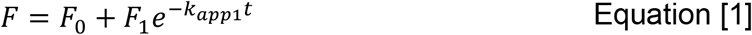

With *t* being the time; *k_app1_*, the apparent rate constant; *F*, the fluorescence; *F_0_*, the initial fluorescence value; and *F_1_*, the fluorescence change amplitude related to *k_app1_*.

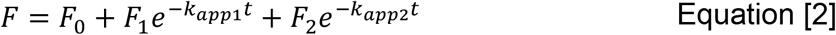

With *t* being the time; *k_app1_*, an apparent rate constant for a first phase; *k_app2_*, an apparent rate constant for a second phase; *F*, the fluorescence; *F_0_*, the initial fluorescence value; *F_1_*, the fluorescence change amplitude related to *k_app1_*; and *F_2_*, the fluorescence change amplitude related to *k_app1_*

30S, IF2, IF1, and mRNA showed conformational changes of IF3_DL_ slower than their respective bimolecular reactions [4], allowing to fit the obtained rates as a function of titrant concentration with a hyperbolic equation (equation 3) where *V*_max_ and *K*_S_ could be calculated.

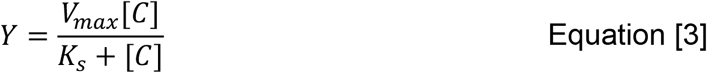

With *Y* being the reaction rate; [*C*], the concentration; *Vmax*, the maximum rate; and Ks, the rapid equilibration constant.

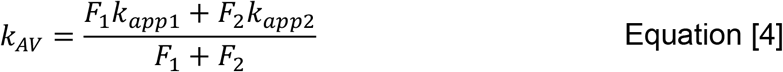

With *k_AV_* being the averaged apparent rate constant calculated from equation 2. Standard errors of *k_AV_* were calculated according to error propagation functions for addition and product using the mean and standard deviation of each term of equation 4.

### Structural modelling

Structures of the 30S pre-IC and 30S IC were obtained from the Protein Data Bank (PDB s: 5lmn, 5lms, 5lmt, 5lmu, 5lmv) [20]. IF3 was extracted into new PDB files from each complex using Chimera software [32]. IF3E166C was then modelled with the Swiss Modeler server using each IF3 pdb file as template [33]. Finally, the modeled IF3E166C replaced the original IF3 in the reported 30S IC structures. fMet-tRNA^fMet^ was removed from all PDB files to generate a subset of structures lacking the tRNA. The accessible volume (AV) of each fluorophore on the 30S ICs structures was determined with FRET-restrained Positioning and Screening software [34]. The fluorophore dimension parameters were: linker length: 15 Å; Width: 4.5 Å; Dye radius: 4.5 Å as defined from a model generated with Maestro software (Schrödinger, NY, USA). The distances between points of the AV of donor and acceptor dyes were calculated in Matlab (Mathworks, USA) using an analytical geometry approximation. The distance distribution between AVs was determined with Prism 7.0 (Graphpad Software, USA).

## Acknowledgments

We specially thank Anna Maria Giuliodori for critically reading the manuscript. This work was supported by InnóvatePerú grants 382-PNICP-PIBA-2014 and 297-INNOVATEPERU-EC-2016 (to PM), by the Fondecyt grant 154-2017-Fondecyt (to PM), and by FIRB Futuro in Ricerca grant (RBFR130VS5 001) from the Italian Ministero dell’Istruzione, dell’Universitá e della Ricerca (to AF). Part of the work on structural dynamics of the ribosome was supported by Russian Science Foundation Grant 17-14-01416 (to ALK).

## Author Contributions

AF and PM conceived the project; JAN and AF performed experiments; JAN, AK, RS, AF, and PM analyzed the data; AF, RS and PM wrote the manuscript with the inputs of JAN and AK.

## Supplementary Information

**Supplemental Figure 1.**
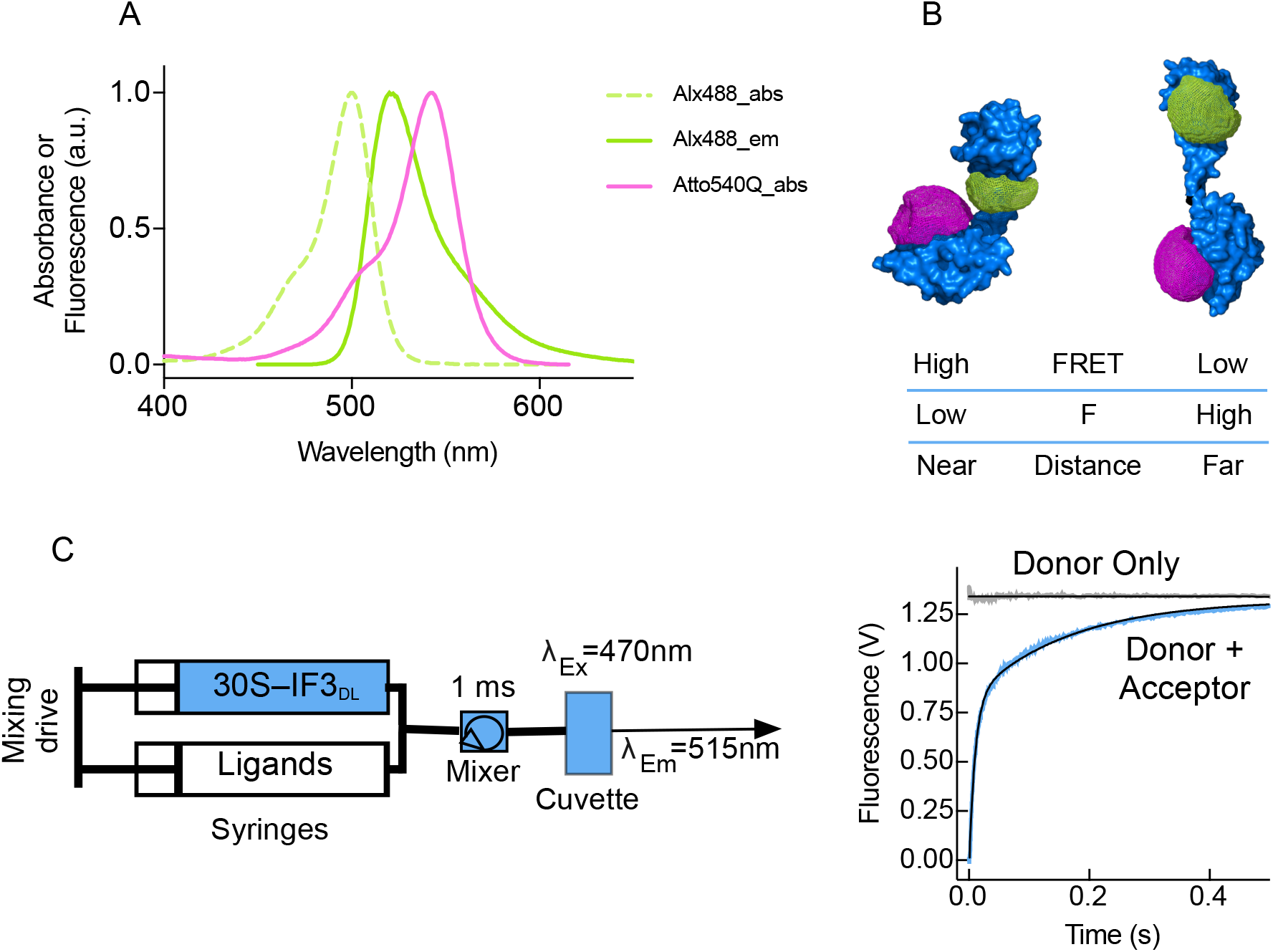
Experimental approach. IF3_DL_ for dynamic measurements of the factor on the 30S platform. **(A)** Absorption and emission spectra of Alexa488 fluorescent dye (green) and Atto540Q quencher (magenta). **(B)** Schematic example of IF3_DL_ arrangements. IF3 surface is shown in blue while meshed volumes indicate the spatial volume that dyes occupy (colors are as (A)). **(D)** Scheme of Stopped-Flow experimental set-up and the typical signal read out upon mixing 30S–IF3_DL_ with a 30S binder (sky blue trace). In order to assign the signal as FRET, the same experiment is performed in the absence of the acceptor, in this case IF3_N Alx488_ (grey trace).

**Supplementary figure 2.**
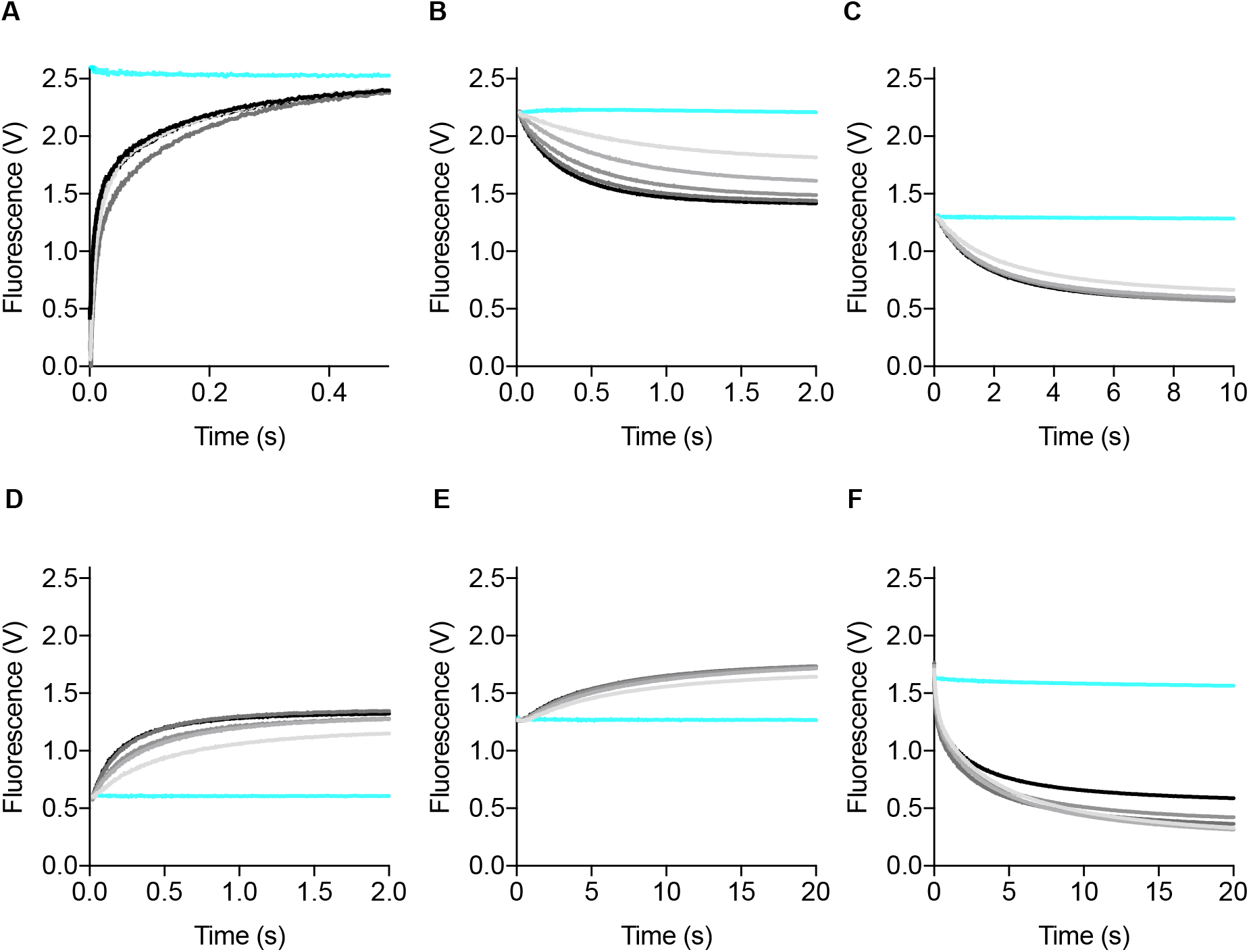
Time traces of IF3_DL_ donor fluorescence change upon formation of all initiation intermediates. **(A)** Time courses of donor fluorescence change of 0.04 μM IF3_DL_ during binding to varying concentration of 30S (0.15-0.5 μM). **(B)** Time courses of IF2 (0.15-0.5 μM) binding to 0.05 μM 30S–IF3_DL_ complexes. **(C)** Time courses of IF1 (0.15-0.5 μM) binding to 0.05 μM 30S–IF2–IF3_DL_ complexes. **(D)** Time courses of fMet-tRNA^fMet^ (0.15-0.5 μM) binding to 0.05 μM 30S–IF3_DL_–IF1–IF2 complexes. **(E)** Time courses of mRNA (0.15-1 μM) binding to 0.05 μM 30S PICs (IF3_DL_). **(F)** Time courses of 50S (0.15-1 μM) binding to 0.05 μM 30SIC (IF3_DL_) complexes. A control against buffer is shown in each titration (sky blue). Each time trace results from the average of 7-10 independent measurement.

**Supplementary figure 3.**
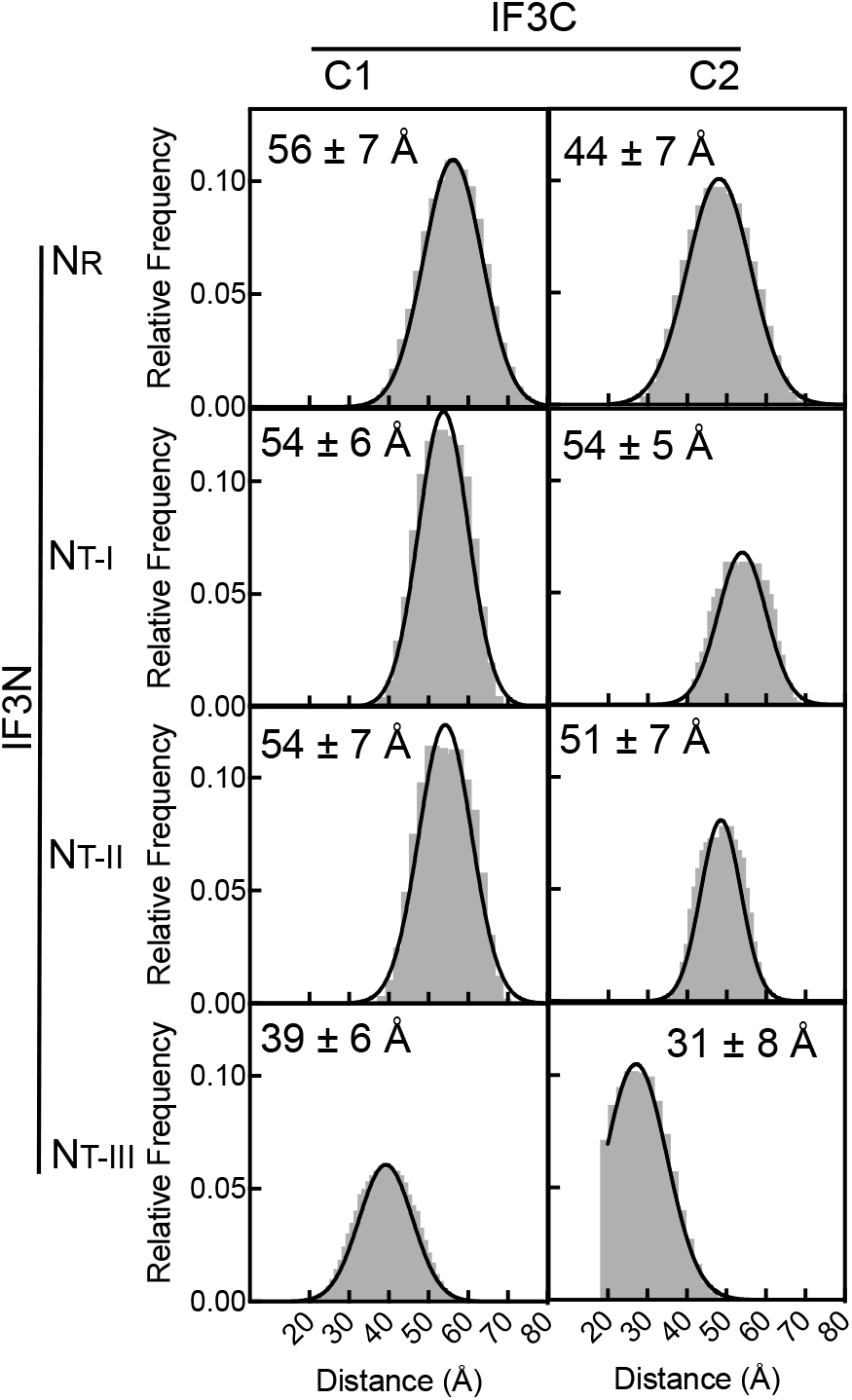
Structural analysis of IF3_DL_ on 30S complexes. Frequency distribution of the distances between donor and acceptor dyes for IF3 occupying all combination of binding sites. The median plus its deviation is indicated.

**Supplementary figure 4.**
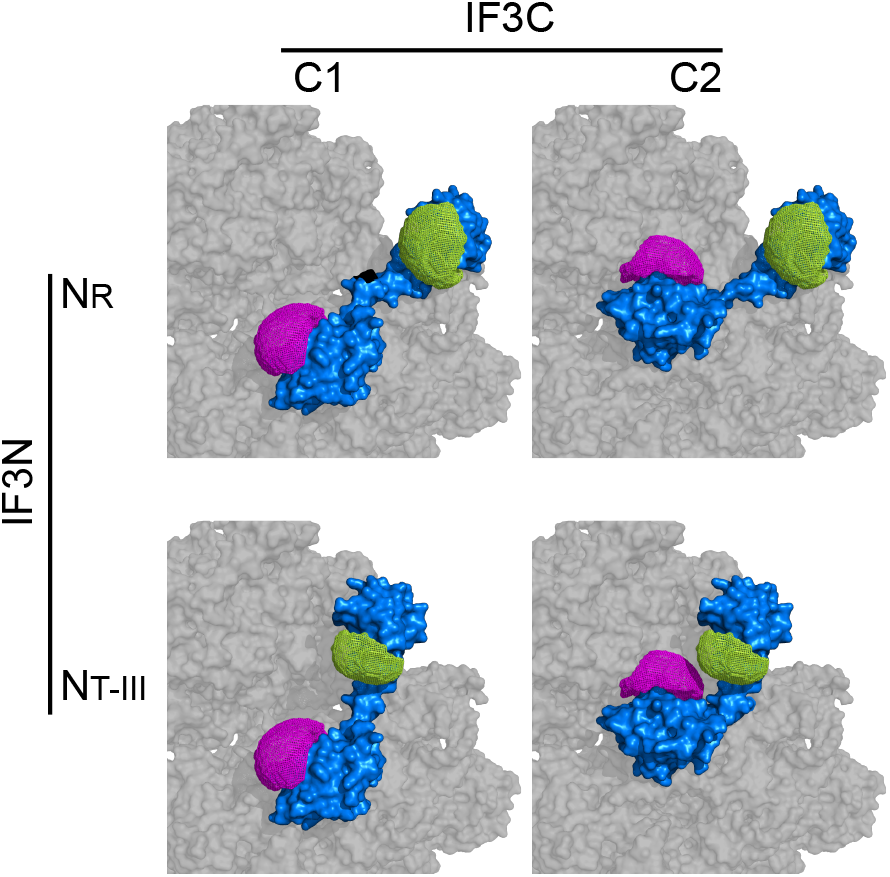
Structural representation of IF3_DL_on 30S complexes. IF3 domains in the Nr, NTIII, C1, and C2 binding configurations in the 30S platform. Donor (Alexa 488) accessible volume (AV) is shown in green and acceptor (Atto540Q) AV in magenta. All structures are redrawn from [20] after modeling IF3 and the dyes attached to the factor (see methods).

**Supplementary Table 1.** Distances between fluorophores and domains on IF3_DL_ layouts. *Distances determined from the AV. **Distance between labelled cysteines in each domain.

| Position | Distance between fluorophores (Å)* | Distance measurements (N x 10<sup>5</sup>) | Distance between domains (Å)** |
| --- | --- | --- | --- |
| C<sub>1</sub> – N<sub>R</sub> | 57 ± 7 | 15.6 | 63 |
| C<sub>2</sub> – N<sub>R</sub> | 44 ± 7 | 14.9 | 61 |
| C<sub>1</sub> – N<sub>T<sub>I</sub></sub> | 54 ± 6 | 8.8 | 57 |
| C<sub>2</sub> – N<sub>T<sub>I</sub></sub> | 54 ± 5 | 7.3 | 51 |
| C<sub>1</sub> – N<sub>T<sub>II</sub></sub> | 54 ± 6 | 9.9 | 60 |
| C<sub>2</sub> – N<sub>T<sub>II</sub></sub> | 51 ± 5 | 6.2 | 50 |
| C<sub>1</sub> – N<sub>T<sub>III</sub></sub> | 39 ± 6 | 11.1 | 54 |
| C<sub>2</sub> – N<sub>T<sub>III</sub></sub> | 31 ± 8 | 9.7 | 46 |

**Supplementary Table 2.** Calculated accessible volume (AV) for each fluorophore at all binding configurations of IF3_DL_. AV spatial coordinates were generated with the FPS-software [34]. *Volume approximation by the number of coordinates to the AV to a sphere with a radius of 15 Å. **Maximum number of coordinates present in an AV with the following parameters: Linker length: 15 Å; Width: 4.5 Å; Dye radius: 4.5 Å.

|  | AV pts | Volume (Å<sup>3</sup>)* | AV pts<br>(no tRNA) | Volumen (Å<sup>3</sup>)*<br>(no tRNA) |
| --- | --- | --- | --- | --- |
| C<sub>2</sub> | 4338 | 3805.82 | 4338 | 3805.82 |
| C<sub>1</sub> | 4975 | 4364.68 | 4975 | 4364.68 |
| N<sub>R</sub> | 3384 | 2968.86 | 3384 | 2968.86 |
| N<sub>TI</sub> | 192 | 168.45 | 2341 | 2053.81 |
| N<sub>TII</sub> | 216 | 189.50 | 2436 | 2137.16 |
| N<sub>TIII</sub> | 223 | 195.64 | 2009 | 1762.54 |
| Max | 16114** | 14137.17 | 16114** | 14137.17 |

## References

1. Biou V, Shu F, Ramakrishnan V (1995) X-ray crystallography shows that translational initiation factor IF3 consists of two compact alpha/beta domains linked by an alpha-helix. The EMBO Journal 14: 4056–4064.

2. Garcia C, Fortier PL, Blanquet S, Lallemand JY, Dardel F (1995) 1H and 15N resonance assignments and structure of the N-terminal domain of Escherichia coli initiation factor 3. Eur J Biochem 228: 395–402.

3. Garcia C, Fortier PL, Blanquet S, Lallemand JY, Dardel F (1995) Solution structure of the ribosome-binding domain of E. coli translation initiation factor IF3. Homology with the U1A protein of the eukaryotic spliceosome. Journal of Molecular Biology 254: 247–259.

4. Milon P, Maracci C, Filonava L, Gualerzi CO, Rodnina MV (2012) Real-time assembly landscape of bacterial 30S translation initiation complex. Nat Struct Mol Biol 19: 609–615.

5. Milon P, Konevega AL, Gualerzi CO, Rodnina MV (2008) Kinetic checkpoint at a late step in translation initiation. Molecular Cell 30: 712–720.

6. Grigoriadou C, Marzi S, Pan D, Gualerzi CO, Cooperman BS (2007) The translational fidelity function of IF3 during transition from the 30 S initiation complex to the 70 S initiation complex. Journal of Molecular Biology 373: 551–561.

7. Antoun A, Pavlov MY, Tenson T, Ehrenberg M MÅ (2004) Ribosome formation from subunits studied by stopped-flow and Rayleigh light scattering. Biol Proced Online 6: 35–54.

8. Gualerzi CO, Brandi L, Caserta E, Garofalo C, Lammi M, La Teana A, Petrelli D, Spurio R, Tomsic J, Pon CL (2001) Initiation factors in the early events of mRNA translation in bacteria. Cold Spring Harb Symp Quant Biol 66: 363–376.

9. Petrelli D, LaTeana A, Garofalo C, Spurio R, Pon CL, Gualerzi CO (2001) Translation initiation factor IF3: two domains, five functions, one mechanism? The EMBO Journal 20: 4560–4569.

10. Ramakrishnan V (2002) Ribosome structure and the mechanism of translation. Cell 108: 557–572.

11. Goyal A, Belardinelli R, Maracci C, Milon P, Rodnina MV (2015) Directional transition from initiation to elongation in bacterial translation. Nucleic Acids Research 43: 10700–10712.

12. Goyal A, Belardinelli R, Rodnina MV (2017) Non-canonical Binding Site for Bacterial Initiation Factor 3 on the Large Ribosomal Subunit. Cell Rep 20: 3113–3122.

13. Pon C, Cannistraro S, Giovane A, Gualerzi CO (1982) Structure-function relationship in Escherichia coli initiation factors. Environment of the Cys residue and evidence for a hydrophobic region in initiation factor IF3 by fluorescence and ESR spectroscopy. Arch Biochem Biophys 217: 47–57.

14. Fabbretti A, Milon P, Giuliodori AM, Gualerzi CO, Pon CL (2007) Real-Time Dynamics of Ribosome-Ligand Interaction by Time-Resolved Chemical Probing Methods. In, Translation Initiation: Reconstituted Systems and Biophysical Methods pp 45–58. Elsevier.

15. Sette M, Spurio R, van Tilborg P, Gualerzi CO, Boelens R (1999) Identification of the ribosome binding sites of translation initiation factor IF3 by multidimensional heteronuclear NMR spectroscopy. RNA 5: 82–92.

16. Moazed D, Samaha RR, Gualerzi CO, Noller HF (1995) Specific protection of 16 S rRNA by translational initiation factors. Journal of Molecular Biology 248: 207–210.

17. McCutcheon JP, Agrawal RK, Philips SM, Grassucci RA, Gerchman SE, Clemons WM, Ramakrishnan V, Frank J (1999) Location of translational initiation factor IF3 on the small ribosomal subunit. Proceedings of the National Academy of Sciences 96: 4301–4306.

18. Dallas A, Noller HF (2001) Interaction of translation initiation factor 3 with the 30S ribosomal subunit. Molecular Cell 8: 855–864.

19. Pioletti M, Schlünzen F, Harms J, Zarivach R, Glühmann M, Avila H, Bashan A, Bartels H, Auerbach T, Jacobi C, et al. (2001) Crystal structures of complexes of the small ribosomal subunit with tetracycline, edeine and IF3. The EMBO Journal 20: 1829–1839.

20. Hussain T, Llácer JL, Wimberly BT, Kieft JS, Ramakrishnan V (2016) Large-Scale Movements of IF3 and tRNA during Bacterial Translation Initiation. Cell 167: 133–144.e13.

21. Julián P, Milon P, Agirrezabala X, Lasso G, Gil D, Rodnina MV, Valle M (2011) The Cryo-EM structure of a complete 30S translation initiation complex from Escherichia coli. PLoS Biol 9: e1001095.

22. López-Alonso JP, Fabbretti A, Kaminishi T, Iturrioz I, Brandi L, Gil-Carton D, Gualerzi CO, Fucini P, Connell SR (2017) Structure of a 30S pre-initiation complex stalled by GE81112 reveals structural parallels in bacterial and eukaryotic protein synthesis initiation pathways. Nucleic Acids Research 45: 2179–2187.

23. Chulluncuy R, Espiche C, Nakamoto JA, Fabbretti A, Milon P (2016) Conformational Response of 30S-bound IF3 to A-Site Binders Streptomycin and Kanamycin. Antibiotics (Basel) 5: 38.

24. Elvekrog MM, Gonzalez RL (2013) Conformational selection of translation initiation factor 3 signals proper substrate selection. Nat Struct Mol Biol 20: 628–633.

25. Liu Q, Fredrick K (2016) Intersubunit Bridges of the Bacterial Ribosome. Journal of Molecular Biology 428: 2146–2164.

26. Antoun A, Pavlov MY, Lovmar M, Ehrenberg M (2006) How Initiation Factors Maximize the Accuracy of tRNA Selection in Initiation of Bacterial Protein Synthesis. Molecular Cell 23: 183–193.

27. Milon P, Carotti M, Konevega AL, Wintermeyer W, Rodnina MV, Gualerzi CO (2010) The ribosome-bound initiation factor 2 recruits initiator tRNA to the 30S initiation complex. EMBO Rep 11: 312–316.

28. Gualerzi CO, Pon CL (2015) Initiation of mRNA translation in bacteria: structural and dynamic aspects. Cell Mol Life Sci 72: 4341–4367.

29. Grigoriadou C, Marzi S, Pan D, Gualerzi CO, Cooperman BS (2007) The translational fidelity function of IF3 during transition from the 30 S initiation complex to the 70 S initiation complex. Journal of Molecular Biology 373: 551–561.

30. Grigoriadou C, Marzi S, Kirillov S, Gualerzi CO, Cooperman BS (2007) A quantitative kinetic scheme for 70 S translation initiation complex formation. Journal of Molecular Biology 373: 562–572.

31. Milon P, Konevega AL, Peske F, Fabbretti A, Gualerzi CO, Rodnina MV (2007) Transient kinetics, fluorescence, and FRET in studies of initiation of translation in bacteria. Meth Enzymol 430: 1–30.

32. Pettersen EF, Goddard TD, Huang CC, Couch GS, Greenblatt DM, Meng EC, Ferrin TE (2004) UCSF Chimera--a visualization system for exploratory research and analysis. J Comput Chem 25: 1605–1612.

33. Waterhouse A, Bertoni M, Bienert S, Studer G, Tauriello G, Gumienny R, Heer FT, de Beer TAP, Rempfer C, Bordoli L, et al. (2018) SWISS-MODEL: homology modelling of protein structures and complexes. Nucleic Acids Research 46: W296–W303.

34. Kalinin S, Peulen T, Sindbert S, Rothwell PJ, Berger S, Restle T, Goody RS, Gohlke H, Seidel CAM (2012) A toolkit and benchmark study for FRET-restrained high- precision structural modeling. Nature Methods 9: 1218–1225.

